# Integrative Computational and Experimental Analysis of Curly Su Mutations in Drosophila melanogaster

**DOI:** 10.1101/2025.09.15.676221

**Authors:** Adebiyi Sobitan, Md Shah Jalal, Qiaobin Yao, Melissa Constantin, Jinrui Xu, Ginger Hunter, Atanu Duttaroy, Shaolei Teng

## Abstract

The Curly Su (dMPO) protein, a homolog of the human myeloperoxidase (hMPO), is critical for wing development in Drosophila melanogaster. Like human peroxidases, dMPO is involved in various cellular and physiological processes, producing significant quantities of reactive oxygen species that contribute to both development and immunity in the fruit fly. Given the significant sequence and structural similarities between dMPO and hMPO, dMPO serves an ideal model for studying peroxidase functions and related pathologies. We performed saturated computational mutagenesis on dMPO, analyzing the effects of 11,191 missense mutations on its stability. Notably, the G378W mutation exhibited the greatest destabilizing effect, while the W621R, potentially pathogenic, also reduced dMPO stability. To investigate these effects *in vivo*, we used genome editing to generate the transgenic Drosophila with G378W, W621R, and deletion of residues 305-687. Remarkably, G378W mutants displayed significant alterations in wing morphology and reduced lifespan. RNA-seq analysis of transgenic and wild-type flies revealed differentially expressed genes (DEGs), as interpreted through gene ontology analysis. Our integrated computational and genetic approach identified dMPO mutations that disrupt protein stability and alter gene expression. These findings offer new insight into how single-point mutations can lead to systemic biological changes.

**Author Summary:** Proteins are the molecular machines that drive almost every process in living organisms, and even small genetic changes can disrupt their structure and function. In this study, we focused on Curly Su, a protein in fruit flies that is similar to human myeloperoxidase. We used a computational approach called saturation mutagenesis to model over 11,000 possible single mutations in the Curly Su protein and predict how they would affect its stability. We then used genome editing to introduce several destabilizing and potential disease-causing mutations into flies. These mutant flies developed abnormal wings, had shorter lifespans, and showed widespread changes in gene activity, particularly in metabolism and immune pathways. By combining computational predictions with experimental validation, our work demonstrates a powerful and generalizable strategy for linking specific genetic mutations to whole-organism outcomes. This approach not only sheds light on the biological role of Curly Su but also provides a framework that can be applied to other genes and organisms, offering insights relevant to human health and disease.

## Introduction

The Curly Su (*drosophila* myeloperoxidase, *d*MPO) is a heme peroxidase that primarily helps *Drosophila melanogaster (Fruit fly)* respond to oxidative stress and wings maturation.

The *d*MPO uses hydrogen peroxide (H_2_O_2_) to crosslink proteins and stabilizes the wing cuticles during wing formation [1]. However, an increase in H_2_O_2_ concentration serves as a signal to initiate the development of the *Curly* phenotype [2]. The *cysu* gene (*dMPO*) is located on chromosome 3R, which encodes the Curly Su protein. The *Curly* phenotype was first observed over 90 years ago, where the wings curve upwards with less rigid venation and fine wrinkles at the end. The function of the *d*MPO is to suppress the *Curly* wing phenotype, hence its name, Curly Suppressor or Curly Su [3]. Heterozygous mutation of *cysu* typically defines the *Curly* phenotype, while homozygous mutation is usually lethal to the fruit fly. The *Curly* phenotype is, therefore, a dominant mutant wing characteristic [4]. The *d*MPO is a highly conserved protein in *drosophila* species and invertebrates. The *d*MPO has a peroxinectin domain with a cell adhesive property, which shares 51% sequence similarities with a peroxinectin protein discovered in shrimp [5] [6]. Northern blot analysis showed that peroxinectin might play a role in the host defense mechanism [6]. Myeloperoxidase in humans (*h*MPO) primarily contributes to the immune system by destroying foreign organisms. Like *d*MPO, *h*MPO utilizes H_2_0_2_ to create super-oxides toxic against foreign pathogens [7].

The fruit fly is an effective animal model with conserved genetics. Researchers can maintain different strains of fruit flies for a long period of time, which can be used in the future to address complex genetic problems. In recent years, the use of fruit fly has increased in biomedical science and developmental biology research [8]. For instance, a Nobel-prized-research discovered that most fruit fly genes responsible for embryonic development are homologous to genes responsible for human development [9]. Another study used fruit fly to prove the chromosomal theory of inheritance, by exploring the *white* gene on the X chromosomes [10]. Further, fruit fly has been used to also understand genetics by defining the mutational effects of X-rays on genes [11]. The exploration of the fruit fly in drug discovery holds a huge potential in the identification of rare variants in humans [12]. Unlike experiments that are time-consuming, laborious, and expensive, computational approach is a more reliable alternative. We implemented a high-throughput, structure-based mutagenesis workflow that evaluated over 11,000 missense mutations in dMPO, integrating multiple prediction tools and genetic experiments to systematically assess the nutation effects on protein functions. Our research demonstrates the value of integrating computational and experimental approaches to rapidly elucidate the phenotypic effects of novel missense mutations, providing insights into the molecular mechanisms of MPO-related functions and diseases.

## Results

### Drosophila Curly Su (*d*MPO) and human Myeloperoxidase (*h*MPO) proteins share some homology

The sequences of *d*MPO (Q9VEJ9-1) and *h*MPO (P05164-1) optimally aligns with 29.1% (248/851) identity and 43.1% (367/851) similarity using the EBLOSUM62 matrix. Moreover, the alignment between *d*MPO’s peroxinectin domain and *h*MPO’s myeloperoxidase_like domain shows higher identity (39.2%) than the overall protein. This level of sequence similarity indicates that the homologs have very similar structures. PyMol aligns the atoms in the residues in the three-dimension space by considering the alpha carbon atoms. The RMSD of the structural alignment of *d*MPO and *h*MPO (PDB ID: 1CXP) is 0.296 (2518 to 2518 atoms), indicating a more conserved structure. Interestingly, residue G378 in *d*MPO corresponds to G402 in *h*MPO. We reported G402, through *in-silico* analysis, to be critical for the structural stability of *h*MPO [13]. In our recent article [13], we analyzed W643R, which changed the structure of *h*MPO and was present in subjects diagnosed with myeloperoxidase deficiency (MPOD) [14]. Our analysis showed that W643R (ΔΔG = 5.92 kcal/mol) destabilizes *h*MPO [13]. Since W643R in hMPO is a conserved site, we targeted its aligned site W621R in *d*MPO (ΔΔG = 5.12 kcal/mol) with the expectation that it will destabilize the *d*MPO structure as well as with some plausible phenotypic similarity to *h*MPO mutant.

### *In silico* mutagenesis of Curly Su (*d*MPO)

An *in-silico* mutagenesis of the Curly Su (*d*MPO) protein was performed with 11191 non-redundant missense mutations in its 589 residues. Next, we computed the free energy changes (ΔΔG) caused by each missense mutation (Supplementary Table 1). Alanine scanning and saturated mutagenesis of *d*MPO predicts that residues G378, G574, D288, and E269 are critical for stability and function of *d*MPO (Fig 1). G378W significantly increased the ΔG of *d*MPO by 87.74 kcal/mol (Table 1), suggesting this mutation has the most destabilizing effect on the protein.

**Fig 1.**
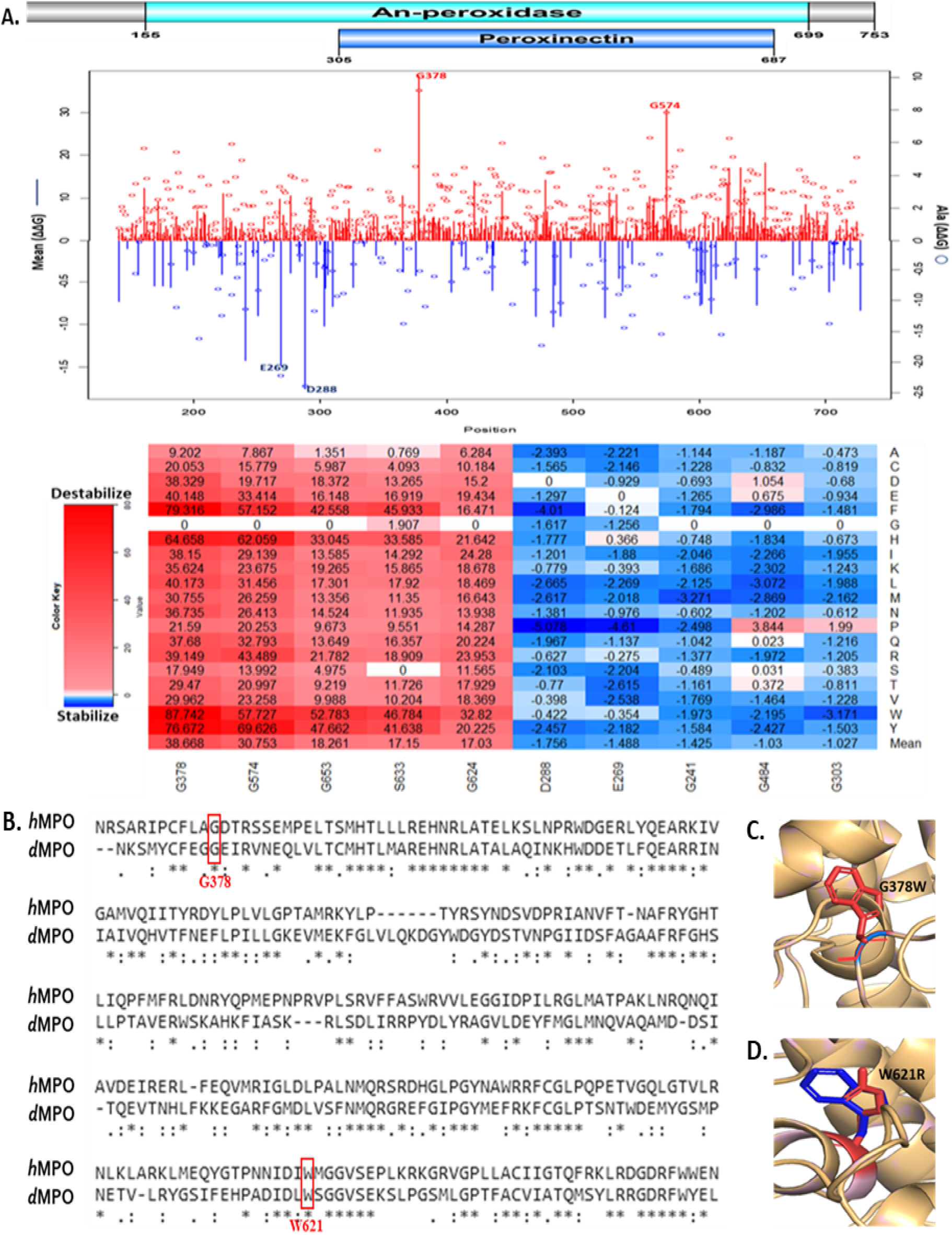
Computational Analysis. **A**. Line chart of the Alanine scanning & saturated mutagenesis of *d*MPO (Top). Heatmap showing 5 most destabilizing and 5 most stabilizing single-site mutagenesis (Bottom). B. Global Pair-wise alignment of *h*MPO and *d*MPO. Conserved residues G378 and W621 in red vertical rectangle. C. PyMol structural alignments showing **C.** G378W and **D.** W621R. Wild type residue in blue and mutant residue in red.

**Table 1.**
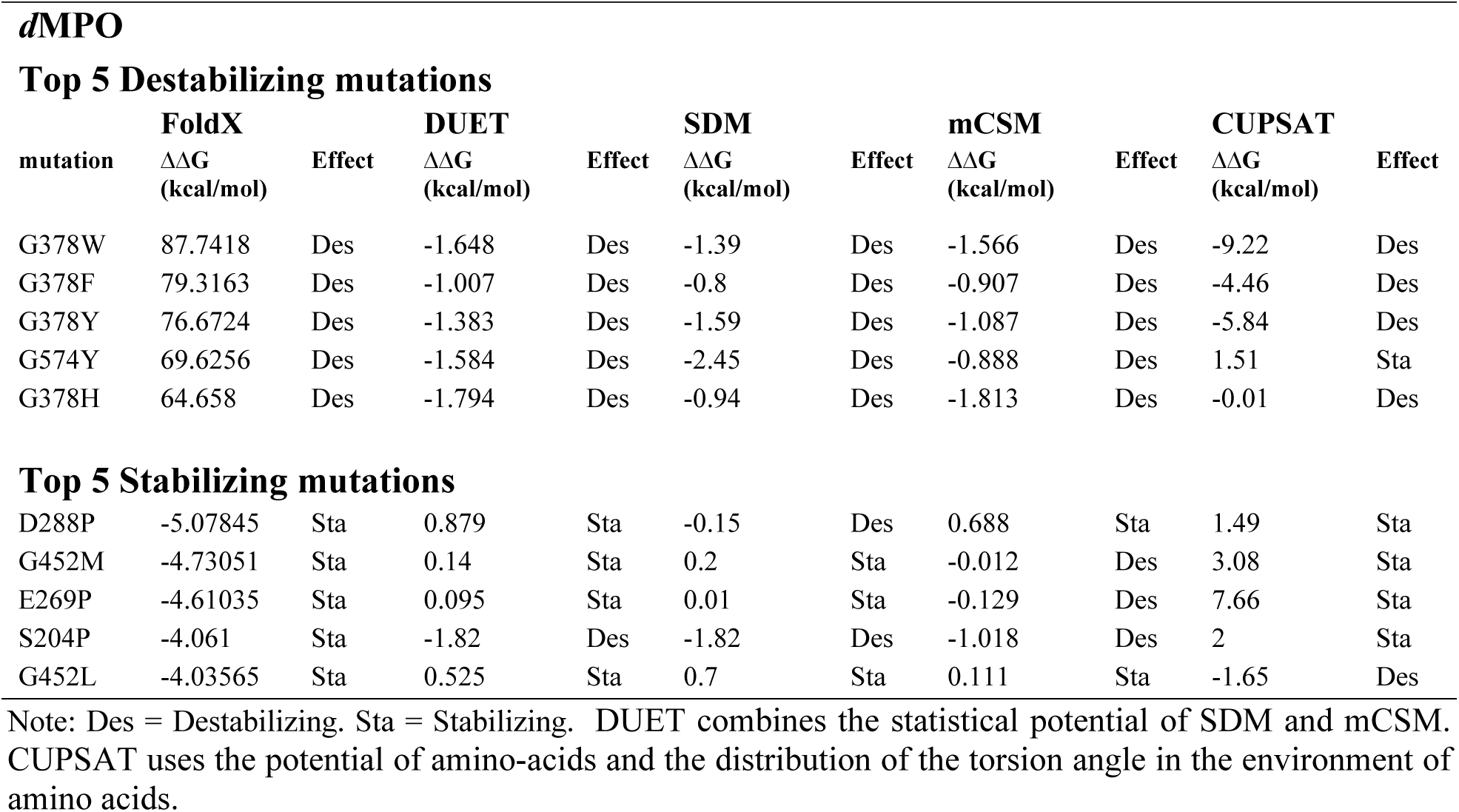
Comparison of Top Missense mutations affecting *d*MPO stability with other computational tools.

**Top 5 Destabilizing mutations**
| mutation | FoldX |  | DUET |  | SDM |  | mCSM |  | CUPSAT |  |
| --- | --- | --- | --- | --- | --- | --- | --- | --- | --- | --- |
| | $\Delta\Delta G$<br>(kcal/mol) | Effect | $\Delta\Delta G$<br>(kcal/mol) | Effect | $\Delta\Delta G$<br>(kcal/mol) | Effect | $\Delta\Delta G$<br>(kcal/mol) | Effect | $\Delta\Delta G$<br>(kcal/mol) | Effect |
| G378W | 87.7418 | Des | -1.648 | Des | -1.39 | Des | -1.566 | Des | -9.22 | Des |
| G378F | 79.3163 | Des | -1.007 | Des | -0.8 | Des | -0.907 | Des | -4.46 | Des |
| G378Y | 76.6724 | Des | -1.383 | Des | -1.59 | Des | -1.087 | Des | -5.84 | Des |
| G574Y | 69.6256 | Des | -1.584 | Des | -2.45 | Des | -0.888 | Des | 1.51 | Sta |
| G378H | 64.658 | Des | -1.794 | Des | -0.94 | Des | -1.813 | Des | -0.01 | Des |
| **Top 5 Stabilizing mutations** |  |  |  |  |  |  |  |  |  |  |
| D288P | -5.07845 | Sta | 0.879 | Sta | -0.15 | Des | 0.688 | Sta | 1.49 | Sta |
| G452M | -4.73051 | Sta | 0.14 | Sta | 0.2 | Sta | -0.012 | Des | 3.08 | Sta |
| E269P | -4.61035 | Sta | 0.095 | Sta | 0.01 | Sta | -0.129 | Des | 7.66 | Sta |
| S204P | -4.061 | Sta | -1.82 | Des | -1.82 | Des | -1.018 | Des | 2 | Sta |
| G452L | -4.03565 | Sta | 0.525 | Sta | 0.7 | Sta | 0.111 | Sta | -1.65 | Des |

Note: Des = Destabilizing. Sta = Stabilizing. DUET combines the statistical potential of SDM and mCSM. CUPSAT uses the potential of amino-acids and the distribution of the torsion angle in the environment of amino acids.

Comparison with other computational tools like SDM, DUET, mCSM, and CUPSAT since all utilize structure-based information to predict ΔΔG values yielded similar consensus among the top 5 destabilizing and top 5 stabilization mutations in Table 1.

### Experimental validation using transgenic Drosophila

Mutations in the *d*MPO gene cause abnormal wing phenotypes and reduced lifespan in adults. Gross RNAi-based knockdown and Minos-based insertion in the *d*MPO gene caused wing deformity in the adult fly [5]. Using CRISPR-Cas9-based point mutagenesis we generated point mutants for W621R, G378W, and we also created a deletion line (Del305-687) spanning both single mutant sites (Fig 2A and 2B). Fruit flies with homozygous *d*MPO mutants G378W and W621R are viable. Similarly, the Del305-687 mutants are also homozygous viable. However, the adult animals for each of these mutant lines show severe wing defects, including bent, shrunken, slender, and fragile wings.

**Fig 2.**
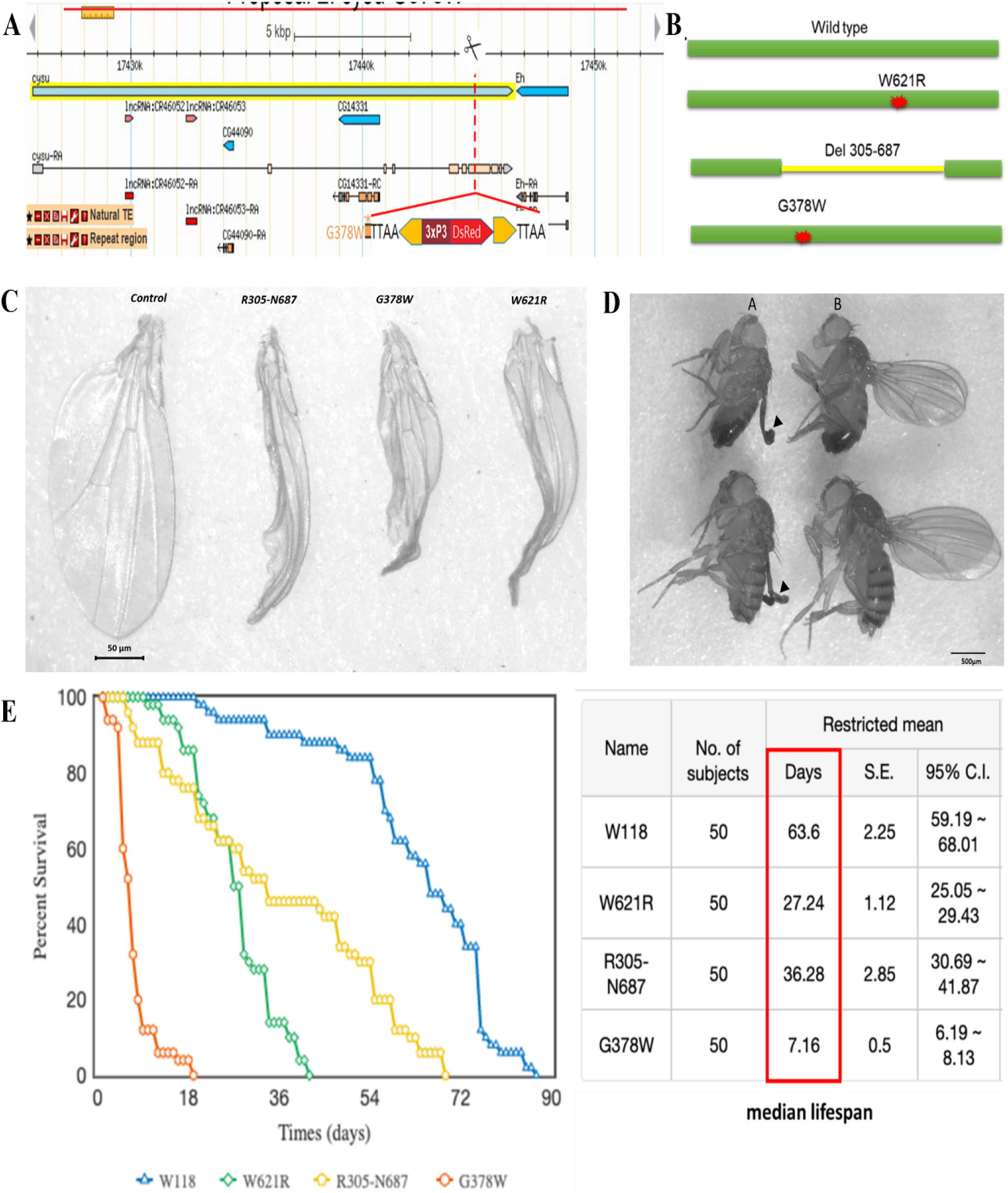
A. PBacDsRed system for transgenesis. B. Schematic of location of the missense mutations and deletion. B. C. Comparison of adult wings in the control and mutant fruit fly strains. D. Homozygous knockdown fly model. Panel A: Homozygous w;; dMPO G378W [CRISPR(PBacDsRed)] male and female. Panel B: Control male and female (w;; dMPO G378W [CRISPR(PBacDsRed)]/TM6B,Tb). Black Arrowhead indicates mutant wing phenotype. E. Survival rate (left) and median lifespan (right) of fruit fly strains.

Immediately after eclosion wings appear like the wild type in all mutants. However, during post eclosion wing formation events, the mutant wings fail to expand, which causes them to appear as slender and shrunken from the dorsal and ventral sides (Fig 2C and 2D). The length of the wing was not affected in any of the mutant lines. Although the newly made mutant lines phenocopies previously reported knockdown of *d*MPO with various ubiquitous and wing-specific drivers, the current study provides new insight into the specific site of the *d*MPO which caused the defect. All three mutants W621R, G378W, and Del305-687 are rescued to complete wild type wings with a dMPO overexpression [5], proving the wing defect appear due to specific mutation at the specific sites of the *d*MPO gene.

### Humanized *MPO* mutants in Drosophila severely reduce life span

While the phenotypic data clearly indicates the role of *d*MPO in wing maturation process, it can’t be the sole function of this protein given the capacity of the enzyme to produce H_2_O_2_. So, we decided to measure the life span of *d*MPO mutant flies which will suggest the impact of the enzyme function in relation to other physiological processes. Therefore, we measured the female life span of three mutant lines (W621R, R305-N687, and G378W) and compare it with a background matched control line (w1118). The longevity of all the *d*MPO mutant lines significantly reduced compared to the control line (Fig 2E). The wild-type flies exhibited a median lifespan of 63 days. In contrast, all the *d*MPO mutant lines, including W621R, R305-N687, and G378W observed significantly reduced median lifespans of 27, 36, and 7 days, respectively. The vulnerable wings of the mutant flies affect the overall fitness and flight of the flies, and this phenomenon leads to reduced lifespan.

### Quality control, statistical analysis and visualization of Differentially Expressed Genes

To assess the transcriptomic impact of these mutations, we conducted RNA-seq and differential gene expression analysis in both mutant and wild-type flies. The Quality control of the raw read from the Illumina encoding flagged none of the 424735740 paired sequences as having poor quality, and no adapter content was present. The output from our De Novo assembly showed 45.76 GC %, a total of 59298 ‘genes’, and 99965 transcripts. We denoted the wild type as A, W621R as B, Del 305-687 as C, and G378W as F. Each sample has three biological replicates. FDR < 0.00005 was used as a threshold for creating subsets of differentially expressed genes (DEG) between treatment pairs. The DEGs are overlayed as parallel coordinate lines on boxplots of each replicate per treatment group (Fig 3). The A – B treatment group has approximately a total of 235 DEGs, A - C has 425 DEGs, and A – F has 495 DEGs. Hierarchical clustering places each DEG into distinct cluster of either downregulated or upregulated genes. Cluster 1 genes in blue are downregulated and Cluster 2 genes in green are upregulated.

**Fig 3.**
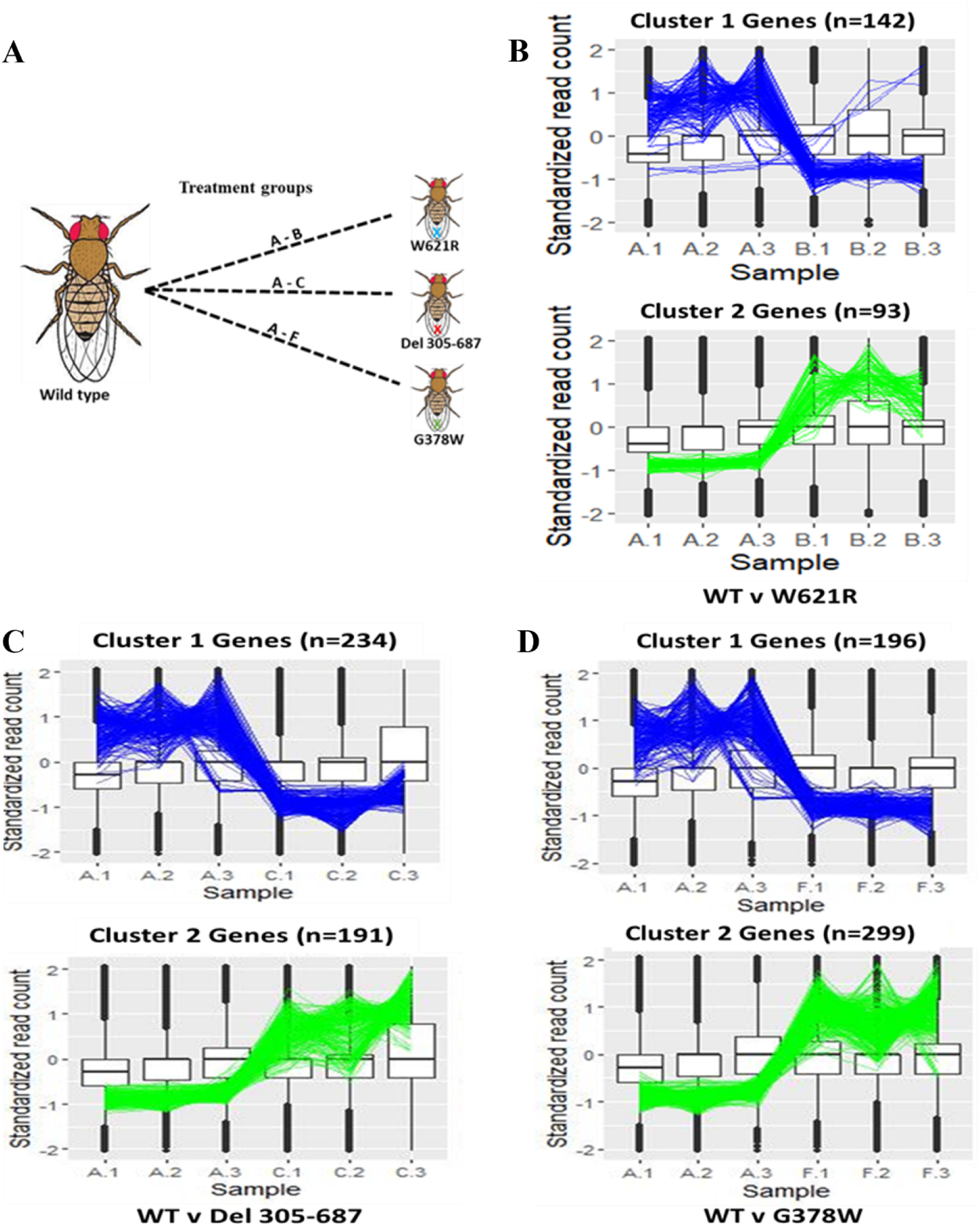
**(A**) Treatment groups of Fruit fly samples and Hierarchical Clustering of DEGs in **(B)** WT v W621R (**C)** WT v Deletion **(D**) WT v G378W.

The scatter plots on the left panel below have overlays of DEGS and display their variability within a sample and between treatment groups (Fig 4). Within a sample, DEGs aggregate around the regression line. However, DEGs either skew to the right (downregulated) or left (upregulated) of the regression line between treatment groups. The volcano plots show the DEGs overlaying the entire genes, using the logarithm of the fold change (logFC) and p-values (-log10(PValue)) as statistical measures. Each hexagonal bin contains genes of varying amounts. For each treatment group, *Adh* and *col* genes were annotated to pinpoint their locations.

**Fig 4.**
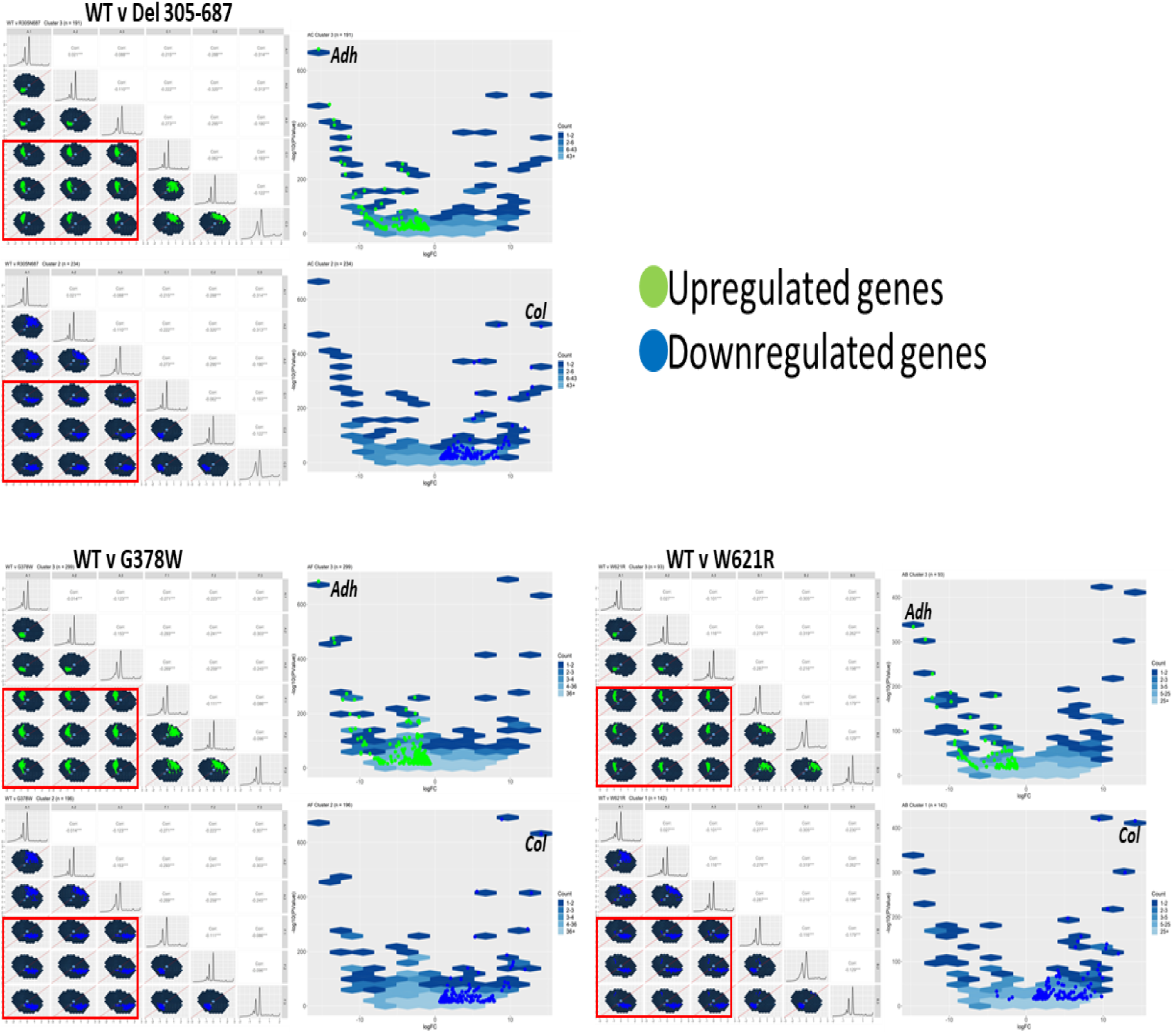
**Overlay of upregulated genes in green and downregulated genes in blue**. The Red rectangles in Scatter plots show the nine treatment groups. The Volcano plots are annotated with *Adh* and *Col* genes.

Table 2 below show the statistical values of the top five upregulated and downregulated DEGs in all treatment groups. Notably, Alcohol dehydrogenase, *Adh*, is the most upregulated gene and cytochrome c oxidase, *col*, is the most downregulated gene in all treatment groups. The logarithmic scale of the fold change (FC) highlights the significance in the expression of genes in the treatment groups.

**Table 2.**
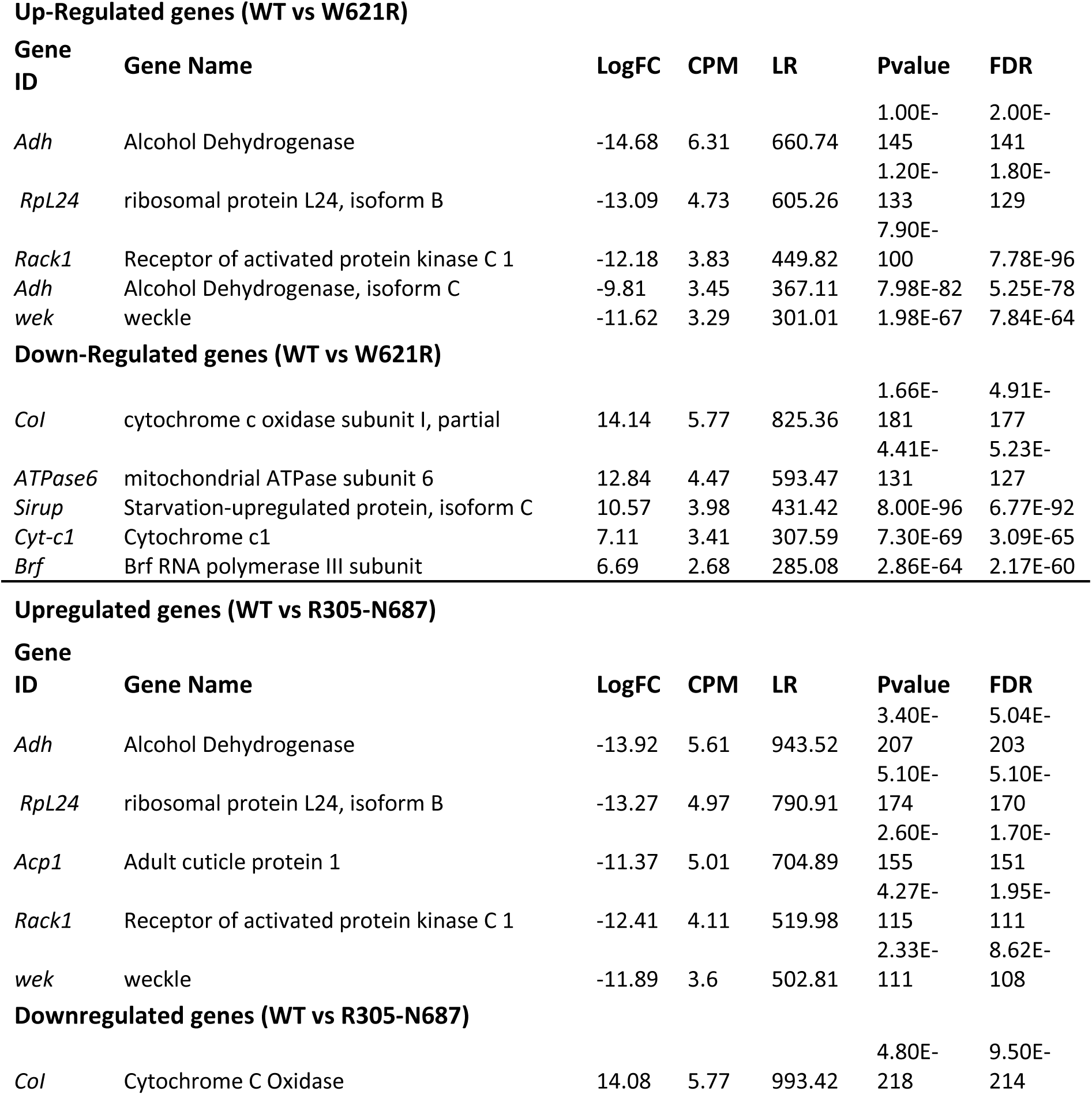
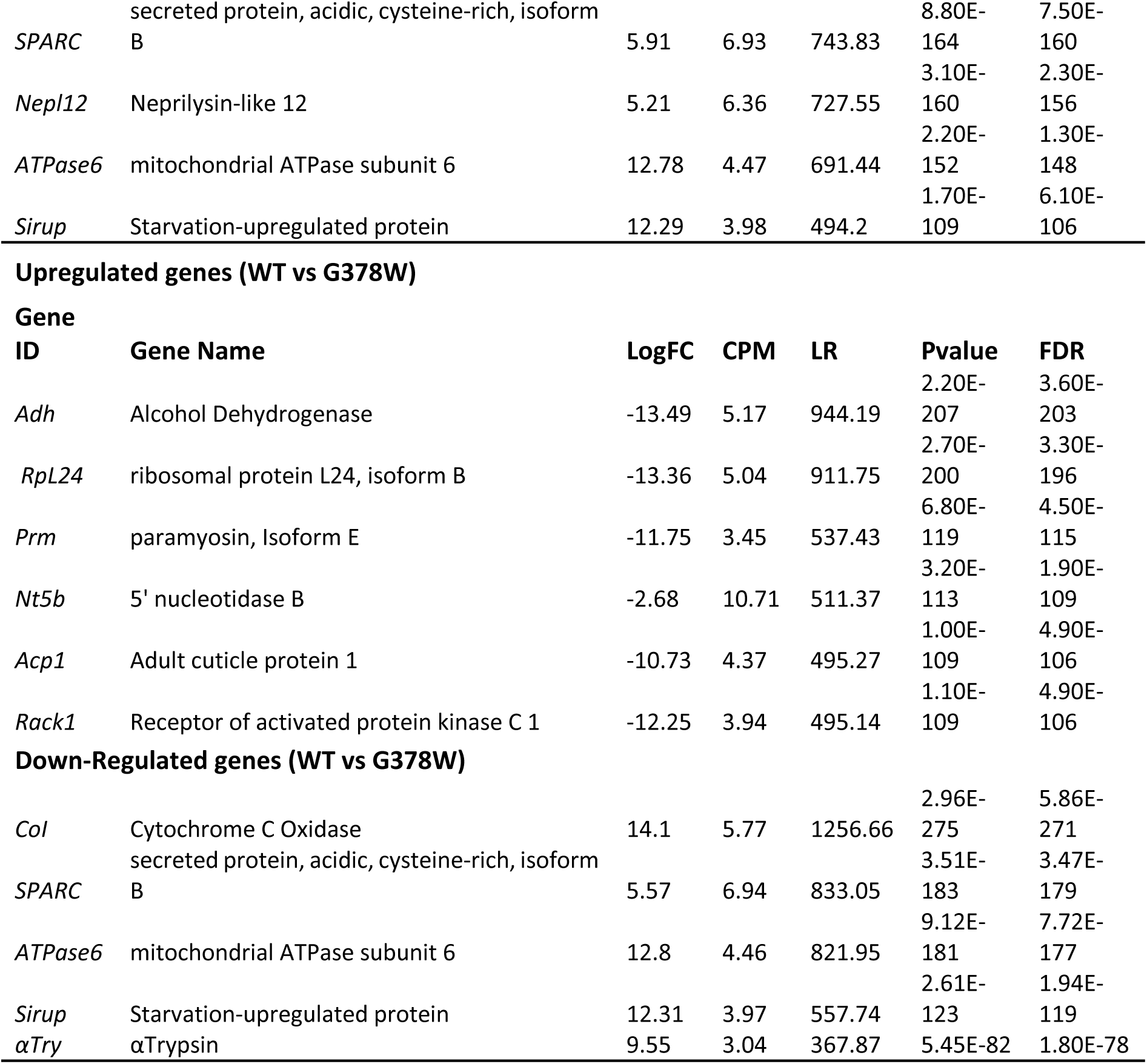
Top upregulated and downregulated genes in fruit fly strains with W621R mutation, deleted Peroxinectin <u>domain (R305-N687), and G378W mutation.</u>.

### Gene Ontology & RT-qPCR of G378W

For both downregulated and upregulated genes in all treatment groups, we annotated subsets of the DEGs (Fig 5). The Venn diagram shows the subsets of annotated genes in each treatment group, and the number of overlapping genes [15]. In fruit flies, the missense mutation W621R of the *d*MPO gene significantly enhances defense response and reduces the glycogen biosynthetic process. The statistically significant GO Terms are shown below, with p-values less than 0.05. Deletion of the Peroxinectin domain (305-687) in the *dMPO* gene significantly increases the overall defense response of the fruit fly while the development of wings is greatly reduced. The computational destabilizing missense mutation of *d*MPO, G378W, evidently increased the response of the fruit fly to bacteria and fungi. Further, G378W upregulates genes whose functions are to mediate innate and humoral immunity in the fruit fly. In contrast, G378W downregulates interspecies interactions and the pathway to glycogen metabolism.

**Fig 5.**
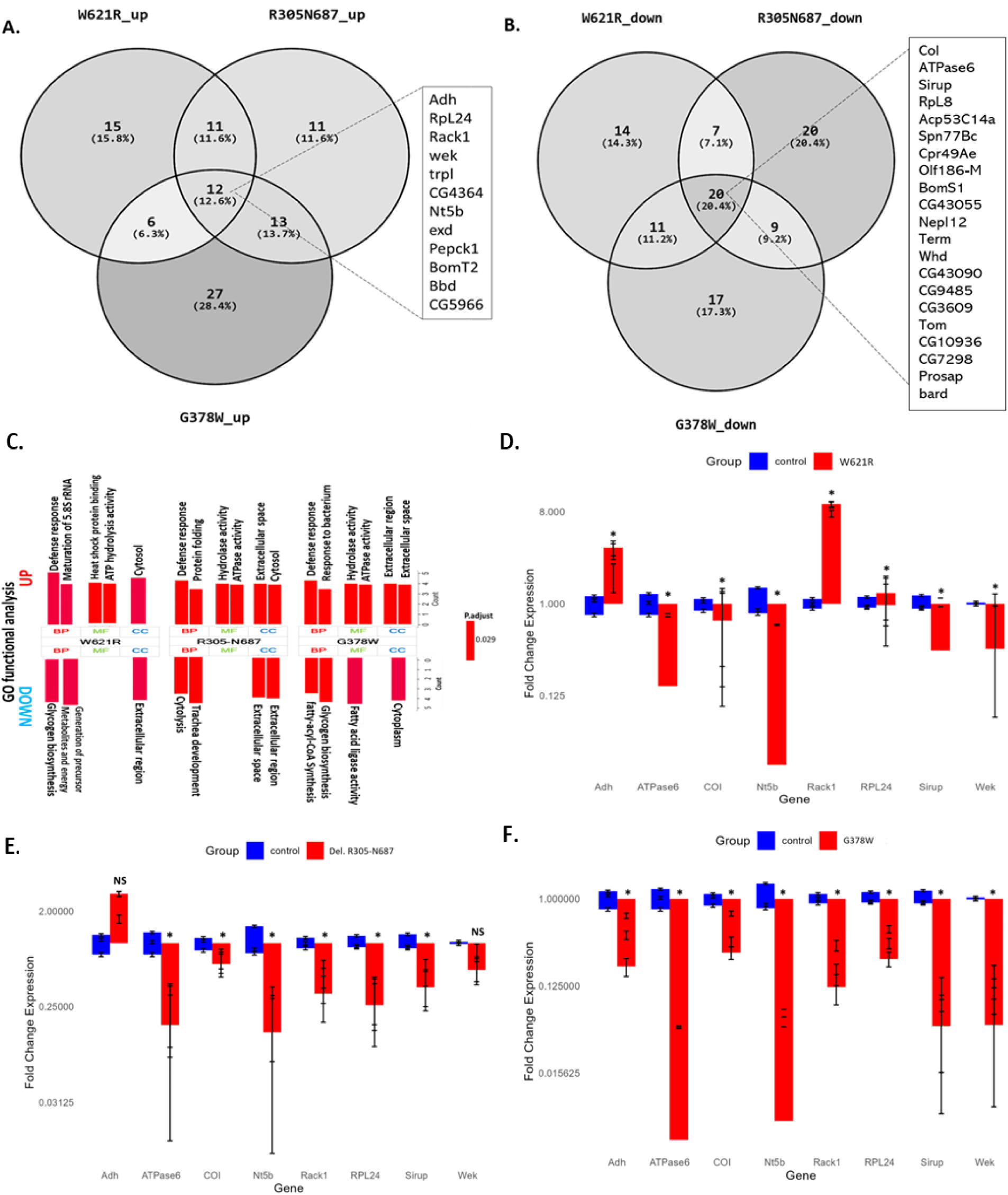
Analysis of Top overlapping DEGs in all three treatment fruit flies. Venn diagram showing the number of overlapping **(A)** Upregulated and **(B)** Downregulated genes in two or more treatment groups. Vertical rectangles show genes common to all three treatment groups. **(C)** GO-Terms of differentially expressed genes (DEGs) in the W621R, R305-N687, and G378W mutant flies. BP-Biological Processes. MF-Molecular Functions. CC-Cellular Components. RT-qPCR of key differentially expressed genes in; **(D)** W621R mutant fruit flies. **(E)** R305-N687 mutant fruit flies, and **(F)** G378W mutant fruit flies. * p-value < 0.05. **^NS^** p-value > 0.05.

To further validate our findings, we performed an RT-qPCR to quantify the variation of gene expressions in the wild-type fruit flies and the G378W mutant fruit flies. The relative fold expression of certain genes in the qPCR experiment coincided with the computational analysis outcomes, while others were either deemed insignificant or showed reversed regulatory pattern. Gene expression was considered statistically significant if the p-value was less than 0.05. Alcohol dehydrogenase (*Adh*) exhibited significant upregulation in the qPCR, as supported by the bioinformatics analysis of DEGs. However, cytochrome c oxidase (*Col*) did not show significant downregulation in the qPCR analysis despite displaying an upregulated pattern.

## Discussion

The *d*MPO is a homolog to the *h*MPO, sharing high sequence and structural similarities. In the *d*MPO, 66% of the missense mutations have ΔΔG greater than 0.5 kcal/mol. Moreover, we observed an increase in the number of destabilizing missense mutations of residues within the peroxinectin domain. The residues in the peroxinectin domain play critical roles in maintaining the stability of *d*MPO. G378W has the highest destabilizing effect on *d*MPO stability. Notably, G378 in dMPO aligns with G402 in hMPO, highlighting potential functional conservation. Although not previously reported, G402 and G378 must play a role in conserving the evolutionary function of *h*MPO and *d*MPO, respectively. The top five stabilizing missense mutations of *d*MPO, D288P, G452M/L, E269P, and S204 correspond to residues P311, L476, P293, and A227, respectively, in the *h*MPO. However, there is no specific information regarding post-translational modifications (PTMs) or mutations at residues P311, L476, P293, and A227 in *h*MPO. Further experimental studies are necessary to elucidate the significance of these residues in *h*MPO’s structure and function.

Genetic screening of G378W in a homozygous knockdown fly model further supports the critical function of G378. The fly model with homozygous G378W exhibited a curly wing phenotype, and a few of the flies (12 out of 225) survived into adulthood. The survival rate (5.33%) of the homozygous G378W knockdown fly is less than the classical mendelian ratio. G378 is a critical residue in *d*MPO, which highlights that *d*MPO helps in wing maturation and possibly developing the fly itself. The DEGs from all three mutant fruit flies mostly overlap. We used false discovery rate (FDR) as the default statistical parameter, and the threshold was set to FDR < 0.0005 to maximize potential DEGs and eliminate false positives. Each mutant fruit flies have an average of one hundred DEGs (fifty up-regulated and fifty down-regulated genes), with twelve and twenty overlapping up-regulated and down-regulated DEGs. The DEG profile was consistent with systemic physiological impacts resulting from dMPO destabilization.

Differential expression analysis revealed several key genes consistently impacted across the mutant lines. The expression of alcohol dehydrogenase (*Adh*) is heightened in all three mutant fruit flies, and research indicates its involvement in ethanol metabolism and the enhanced ethanol tolerance observed in the fruit flies [16]. Further, *Adh* is important in the development and survival of fruit flies and an increase in *Adh* is typically an evolutionary response [17]. Conversely, the down regulation of Cytochrome C Oxidase (*CoI*) is indicative of a dying or prematurely aging fruit fly. The *CoI* has been found to play a role in the mitochondrial respiratory activity of fruit flies and loses its structural integrity in a deteriorating condition [18]. The *ATPase6* gene is involved in oxidative phosphorylation, and the synthesizing of ATP is critical for healthy neurons. The downregulation of the *ATPase6* function causes mitochondrial neurodegeneration in drosophila melanogaster [19]. These results suggest that dMPO dysfunction compromises mitochondrial integrity and metabolic balance, contributing to the reduced viability observed in mutant flies.

In addition to metabolic changes, several genes involved in protein synthesis, immune regulation, and stress adaptation were dysregulated. The *RpL8* gene is conserved in drosophila melanogaster, as well as in humans. *Rpl8* encodes a ribosomal protein critical for protein synthesis—downregulation of *RPL8* via RNA interference results in decreased cell size during eye development in drosophila melanogaster [20]. The withered (*whd*) gene encodes carnitine palmitoyltransferase I (CPT I), responsible for the β-oxidation of long-chain fatty acids in the mitochondria. Mutations or downregulation of the function of *whd* confer hypersensitivity to oxidative stress, increased sensitivity to starvation, and heavy metal toxicity [21]. *Rack1* defends drosophila melanogaster in the event of viral infection. However, *Rack1* upregulation quickly depletes glycogen stores and ultimately impairs autophagic responses [22]. The *wek* gene plays a role in the dorsoventral patterning of the Drosophila embryo. Wek might also have a downstream impact on the immune response in Drosophila. An upregulation of *wek* can cause self-destruction of the drosophila cells [23]. *RPL24* encodes a ribosomal protein, which promotes immune response in the Drosophila. An upregulation of *RPL24* disrupts ribosome synthesis, resulting in altered cell growth and increased cellular stress [24]. The *trpl* gene mediates the phototransduction of the visual system of Drosophila melanogaster. However, excessive upregulation may cause photoreceptor degeneration due to calcium toxicity [25]. Collectively, these gene expression changes reflect a broad network of downstream disruptions that follow from dMPO destabilization and highlight its systemic importance beyond wing development.

## Conclusion

Our findings established *d*MPO as a homolog of *h*MPO, exhibiting significant sequence and structural similarities. The destabilizing impact of missense mutations, particularly within the peroxinectin domain, underscores the domain’s critical role in maintaining *d*MPO stability. Notably, G378W exerts the most profound destabilizing effect, aligning with *h*MPO’s G402, suggesting an evolutionary conserved function. Genetic screening further supports the significance of G378, as homozygous G378W knockdown flies exhibited severe developmental defects and diminished survival, reinforcing *d*MPO’s role in wing maturation and overall fly viability.

Transcriptomic analysis of mutant flies reveals overlapping differentially expressed genes (DEGs) involved in metabolism, mitochondrial function, immune response, and neurodevelopment. The upregulation of *Adh* suggests an adaptive response to environmental stressors, while the downregulation of *CoI* and *ATPase6* indicates mitochondrial dysfunction and neurodegeneration. Additionally, dysregulation of genes such as *RPL8*, *whd*, *Rack1*, *wek*, *RPL24*, and *trpl* reveals broad systemic consequences, including impaired protein synthesis, oxidative stress sensitivity, immune dysregulation, and potential neurotoxicity. While the upregulation of some genes can confer certain advantages, such as improved immune responses or heightened sensory perception, it also carries risks, including metabolic imbalances, cellular stress, or tissue damage.

Together, our results highlight the functional and structural importance of *d*MPO, particularly G378, in both protein stability and organismal development. By linking predicted protein destabilization to transcriptomic changes, our study bridges structural bioinformatics with gene regulation, offering mechanistic insight into how specific missense mutations can reshape gene expression programs. Further experimental validation is needed to elucidate the precise mechanistic roles of homologous residues in *h*MPO and their implications for human health and disease.

## Methods

### Saturated computational mutagenesis

We modeled the structure of *d*MPO, cysu.pdb, using the SWISS-MODEL via the Expasy web server [26].We applied a Perl programming script that substitutes each residue on the *dMPO* structure to 19 other residues, to create a list of all possible missense mutations. The list is saved as “individual_list.txt” and served as an input in the FoldX software suite [27].

Each row in the list has a reference residue, a chain ID, a residue position, a mutant residue, and a semi-colon. FoldX predicts folding energy change (ΔΔG) caused by mutations [28]. The folding energy (ΔG) of the wildtype structure is contributed by the Van der Waals of all atoms (vdw), solvation energy of apolar (solvH) and polar groups (solvP), intra-molecular hydrogen bond (hbond), electrostatic energy (el), backbone entropy cost (Mc), side chain entropy cost (Sc), electrostatic interactions (Kon), extra stabilizing free energy from water molecules (wb), and loss of translational and rotational entropy (Str).

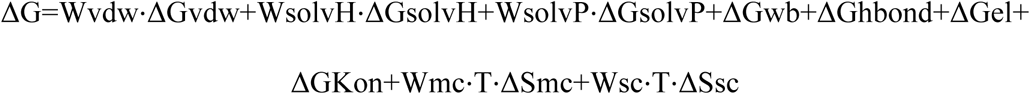

The default energy terms were experimentally determined using protein engineering techniques [29]. FoldX is executed from the command line interface. The Initial step in FoldX mutagenesis is to repair the wild-type structure using the ‘RepairPDB’ command.

Command line example:

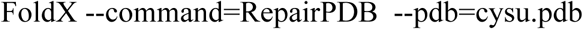

The ‘RepairPDB’ command mutates residues to themselves to attain a low energy conformation. The new structure (1CXP_Repair.pdb) generated by the ‘RepairPDB’ command is used as the wild-type structure in the next step of mutagenesis. Afterwards, we used the ‘BuildModel’ command to calculate changes in folding energies of the mutant structures.

Command line example:

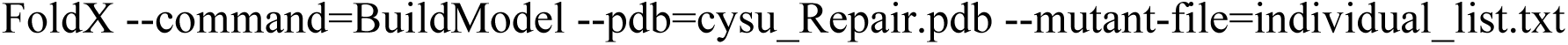

A significant increase or decrease in the mutant structure’s folding energy (ΔG) destabilizes or stabilizes, leading to a less stable or more stable protein, respectively. The change in folding energy (ΔΔG) caused by each mutation can be calculated using the equation:

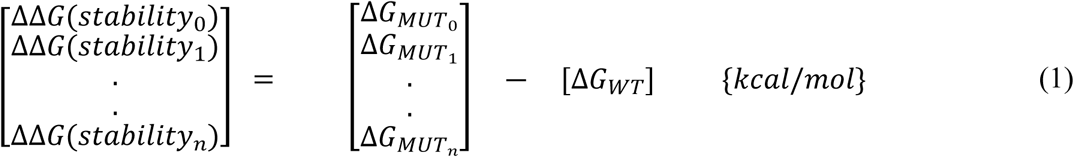

FoldX calculated ΔΔGs deviate from experimental ΔΔGs by 0.46kcal/mol [28]. Therefore, we binned ΔΔG into five categories, in multiples of 0.5 kcal/mol: - highly stabilizing (ΔΔG <-2.0 kcal/mol), stabilizing (-2.0 < ΔΔG <-0.5 kcal/mol), neutral (0.5 < ΔΔG < +0.5 kcal/mol), destabilizing (+0.5 < ΔΔG < 2.0 kcal/mol), and highly destabilizing (ΔΔG > 2.0 kcal/mol).

### Fruit fly strains and husbandry

All experimental and control *Drosophila* strains were maintained on standard cornmeal agar to nurture flies at a constant temperature (23℃ ± 1℃) and on a 12-hour alteration of light/dark cycle. Following *Drosophila melanogaster* stock was obtained from Bloomington Drosophila Stock Centre (BDSC): w1118 (w[1118]).

The following mutant strains were generated using the PBacDsRed system: w[*];; *cysu* R305-N687deletion CRISPR(PBacDsRed)/TM6B, Tb[1], w[*];; *cysu* G378W CRISPR(PBacDsRed)/TM6B, Tb[1], w[*];;cysu W621R CRISPR (inverted PBacDsRed)/TM6B, Tb[1].

The PBacDsRed system facilitated the genetic screening of candidate missense mutations. The selection marker 3xP3-DsRed contains Piggy Bac 3’ terminal repeats, the artificial 3xP3 promoter (three tandem copies of the Pax-6 homodimer binding site and TATA-homology of hsp70), DsRed2, SV40 3’UTR, and Piggy Bac 5’ terminal repeats. It facilitated the genetic screening and then excised by the Piggy Bac transposase. Only one TTAA motif was left after transposition, and after the selection marker was excised.

### Life span Analysis

Life span experiments were performed on standard cornmeal agar medium with at least three biological replicates at 24 ± 1 °C. Newly eclosed female flies from appropriate genotypes were collected immediately and mated with wild-type males for approximately three days. Subsequently, these flies were transferred to vials at a density of 10 flies (5 males and 5 females per vial) and changed the vials every two days to provide fresh food. The quality of the food was strictly monitored/maintained, and the number of dead females was recorded daily. The mortality counts were analyzed using the Online Application for Survival Analysis-2 (OASIS 2) platform using the Log-rank and Wilcoxon tests.

### RNA sample preparation and Sequencing

Adult male and female flies were used from *cysu* mutant and control lines for RNAseq analysis. Newly emerged male and female adult flies (1 day old) were collected from the desired genotype (>100mg) and immediately froze them on dry ice and kept at-80℃ until shipped to Novogene for RNA sequencing. We used three biological replicates for each sample.

RNA samples were purified, sequenced using Illumina NovaSeq 6000 platform, and checked for quality by Novogene (www.novogene.com). The library was checked with Qubit and real-time PCR for quantification and bioanalyzer for size distribution detection.

The output from sequencing the RNAs of our fruit flies was stored in a FastQ format. Each sample has paired reads represented by left and right reads. A total of four paired samples; Wild-type, W621R, deletion R305-N687, and G378W.

### Raw RNA-Seq data preparation and De-Novo Assembly

The De Novo assembly of raw reads produced the “*Trinity.fasta” file for referencing. The new *Trinity.fasta file contains the sequences of all assembled transcripts, which are used for the quantification of genes and transcripts across all four samples. Preparation began with the use of FASTQC to perform quality control on the raw RNA-seq data for each sample. FASTQC screened for the presence of over-represented sequences, the presence of adapters, and the overall base quality of the reads. Then, we used the Trinity software package to assemble the raw RNA-seq data without a reference genome [30]. Trinity performed this task by passing the raw data sequentially through the Inchworm, Chrysalis, and Butterfly modules. RNA-Seq by Expectation-Maximization (RSEM v1.3.3) software suite was used to measure the abundance of transcripts and isoforms or genes across treatment groups [31].

### Statistical analysis of differentially expressed genes (DEGs)

A matrix table was generated to compare raw and standardized read counts of each gene from the wild type to each treatment (transgenic) group. Modification of the bigPint package in R allowed us to create dataMetrics objects for each treatment pair. By default, we generated the Pvalues (significance level), FDR (Multiple comparison significance level, logFC (log base 2-fold change), LR (likelihood ratio), and logCPM (log base 2 counts per million).

The Database for Annotation, Visualization, and Integrated Discovery (DAVID) tool takes lists of the official gene symbols of the DEGs as input and outputs their functional annotations [32] [33].

### RNA isolation and Quantitative PCR

Total RNA was isolated from 10 newly emerged adult flies (male and female) using the Quick-RNA Tissue/Insect Microprep Kit (Zymo Research, Cat # R2030) following the manufacturer’s instructions. The extracted RNA was treated with a DNase I recombinant, RNase-free enzyme (MilliporeSigma, Cat. # 4716728001) to avoid any residual DNA. We used the iScript cDNA kit (Bio-Rad, Cat. # 1708890) to synthesize cDNA from RNA samples. The iTaq Universal SYBR Green Supermix (Bio-Rad, Cat. #1725121) was used to run 20μl qPCR reactions with the manufacturer’s reaction mix preparation and thermal cycling protocol. The data analysis of the qPCR was performed using the Bio-Rad CFX Manager software. Each run was performed in triplicates representing three different biological replicates. Quantitative PCR primers were designed using the NCBI primer blast. Target gene expression was normalized to *rp49* RNA levels.

